# Bfimpute: A Bayesian factorization method to recover single-cell RNA sequencing data

**DOI:** 10.1101/2021.02.10.430649

**Authors:** Zi-Hang Wen, Jeremy L. Langsam, Lu Zhang, Wenjun Shen, Xin Zhou

## Abstract

Single-cell RNA-seq (scRNA-seq) offers opportunities to study gene expression of tens of thousands of single cells simultaneously, to investigate cell-to-cell variation, and to reconstruct cell-type-specific gene regulatory networks. Recovering dropout events in a sparse gene expression matrix for scRNA-seq data is a long-standing matrix completion problem. We introduce Bfimpute, a Bayesian factorization imputation algorithm that reconstructs two latent gene and cell matrices to impute final gene expression matrix within each cell group, with or without the aid of cell type labels or bulk data. Bfimpute achieves better accuracy than other six publicly notable scRNA-seq imputation methods on simulated and real scRNA-seq data, as measured by several different evaluation metrics. Bfimpute can also flexibly integrate any gene or cell related information that users provide to increase the performance. Availability: Bfimpute is implemented in R and is freely available at https://github.com/maiziezhoulab/Bfimpute.

## Introduction

Single-cell RNA-seq (scRNA-seq) has been widely used to study genome-wide transcriptomes in single cell resolution. The cellular resolution made possible by scRNA-seq data distinguishes it from bulk RNA-seq and makes it advantageous in investigating cell-to-cell variation [1]. Today, different commercial platforms are available to perform scRNA-seq, including Fluidigm C1, Wafergen ICELL8 and 10X Genomics Chromium. Droplet-based methods via 10X Genomics Chromium can process tens of thousands of cells; microwell-based, microfluidic-based methods via Fluidigm C1 and Wafergen ICELL8 process fewer cells but with a higher sequencing depth. For all these platforms, missing values make up a large proportion of scRNA-seq data, ranging from 40% - 90% in the gene expression count matrix [2, 3, 4, 5, 6]. In scRNA-seq data, this large percentage of missing events is defined as the so-called ‘dropout’ phenomenon [7]. Gene ‘dropout’ means a gene is observed at a moderate expression level in one cell but it is not detected in another cell of the same type. Analyses of scRNA-seq data, including dimensionality reduction, clustering, and Differential Expression (DE) analysis have shown that effective imputations for dropout events improve downstream analyses and assist biological interpretations [8, 9, 10, 11].

To date, several notable imputation methods have been proposed: scImpute [12], DrImpute [13], MAGIC [14], SAVER [15], VIPER [16] and SCRABBLE [17]. scImpute first performs clustering to identify cell subpopulations and further identifies dropout events through a Gamma-Normal mixture model, finally imputes dropout events by a non-negative least squares regression [12]. DrImpute optimizes the step of identifying cell subpopluations to impute dropout events by averaging the imputation from multiple clustering results [13]. MAGIC builds a Markov affinity-based graph for imputation relying on cell to cell interactions [14]. SAVER uses a Bayesian-based model by various prior probability, and alters all gene expression values [15]. VIPER imputes dropout events relying on local neighborhood cells via non-negative sparse regression models [16]. SCRABBLE has been recently introduced to impute dropout events by adopting the bulk RNA-seq data [17]. Even though a lot of efforts have been taken into analyzing and imputing real dropout events, imputation of dropout events is still a difficult problem because of the high dropout rate and complex cellular heterogeneities for different scRNA-seq datasets. Relying on matrix completion to impute missing values is a long-standing question and has been investigated in biological sciences, including gene expression prediction, miRNA–disease, protein-protein interaction [18] etc. Even though similar mathematical models could be applied to different biological problems, to solve matrix completion problem in scRNA-seq (recovering the dropout events), it is crucial to take the features of scRNA-seq into consideration. Most of existing scRNA-seq imputation methods have shown it is advantageous for imputation to borrow and leverage information from similar cells. In recent years, researchers also start to integrate additional gene or cell related information (e.g. bulk data for SCRABBLE) to assist imputation which is important in matrix completion problem.

In this study, we present Bfimpute, a powerful imputation tool for scRNA-seq data that recovers dropout events by factorizing the count matrix into the product of gene-specific and cell-specific feature matrices [19, 20]. Bfimpute uses full Bayesian inference to describe the latent information for genes and cells and carries out a Markov chain Monte Carlo scheme which is able to easily incorporate any gene or cell related information to train the model and perform the imputation [18] (Figure 1). We demonstrate that Bfimpute performs better than the six other notable published imputation methods mentioned above (scImpute, SAVER, VIPER, DrImpute, MAGIC, and SCRABBLE) in both simulated and real scRNA-seq datasets on improving clustering and differential gene expression analyses and recovering gene expression temporal dynamics (pseudotime analysis) [21].

**Fig. 1.**
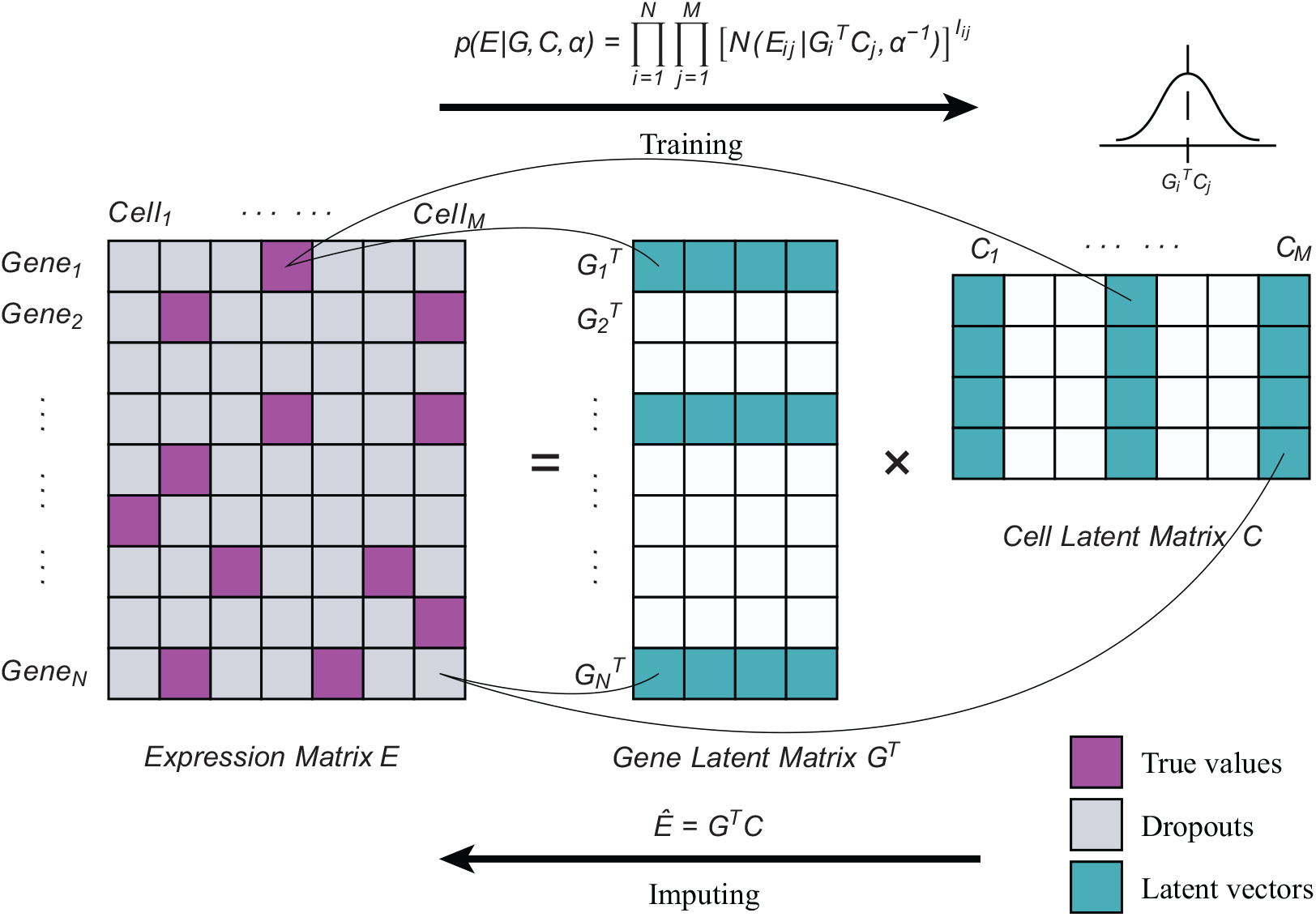
A brief illustration blueprinting the architecture of Bfimpute method. In each group, Bfimpute borrows information from true values and factorizes the expression matrix into two latent matrices using MCMC. After training, Bfimpute imputes dropouts by performing product of the latent matrices. The details are shown in Methods section.

## Methods

### Cell clustering and dropout detection

Bfimpute first provides an optional normalization step to smooth the gene expression values (counts per million, followed by logarithm base 10 with bias 1.01). Bfimpute then performs a local imputation within each cell group. We adopt the same approach as scImpute [12] to detect cell clusters, which applies spectral clustering methods on the result of Principal Component Analysis (PCA) to reduce the impact of dropout events. We integrate spectral clustering by using the ‘Spectrum’ function of the Spectrum R package [22] or the ‘specc’ function of the kernlab R package [23]. Bfimpute also adopts the Gamma-Normal mixture distribution model from scImpute to determine dropout events [12].

### Probabilistic model for scRNA-seq expression matrix imputation

After above-mentioned steps, we adapted a multi-variate priors model from Bayesian Probabilistic Matrix Factorization (BPMF) [20] to recover dropouts for scRNA-seq datasets. Since every cell group is mathematically equivalent, we arbitrarily choose one to demonstrate local imputation in Bfimpute. Suppose we have *N* genes and *M* cells in one cell group, and the expression matrix is *E* ∈ ℝ^*N* ×*M*^. Each entity *E_ij_* represents the expression level of gene *i* in cell *j*. Bfimpute factorizes *E* into *G* ∈ ℝ^*D*×*N*^ and *C* ∈ ℝ^*D*×*M*^ which are defined as gene and cell latent matrix, respectively, where *D* is the dimension of the latent factor. Column vector *G_i_* and *C_j_* represent the gene-specific and cell-specific latent vector, respectively. The imputed matrix to recover *E* will be given as 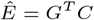.

We introduce the Gaussian noise model for the gene expression profile *E* with precision *α*, which was firstly proposed by Probabilistic Matrix Factorization (PMF) [19]:

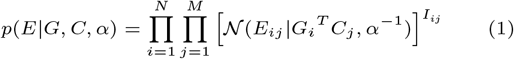

where *I_ij_* is the indicator function that is 0 if the *E_ij_* is a dropout and equal to 1 otherwise.

To get use of gene or cell related information such as bulk data or other data user provided, we add entity features 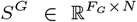 and 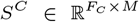 as gene and cell feature matrix, respectively, where *F_G_* and *F_C_* are the dimentionalities of these additional features. The Gaussian model for the prior distributions over genes and cells latent vectors adapted from Macau [18] will be given by:

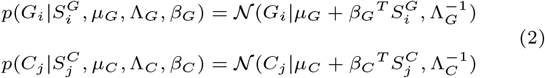

where {*μ_G_, μ_C_*} and {Λ_*G*_, Λ_*C*_} are the means and precisions, and 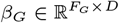 and 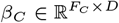 are the weight matrices for the entity features. Weight initialization by a zero mean normal distribution is used and they will be updated iteratively by the Bayesian inference steps (details described later). Also, direct imputation of single cell RNA-seq data could be applied by initiating zeros into feature vectors *S^G^* and *S^C^* (where *F_G_* = *F_C_* = 1) if no additional information is given.

To perform Bayesian inference, we introduce the priors referring to BPMF [20] for {*μ_G_,* Λ_*G*_} and {*μ_C_*, Λ_*C*_}.

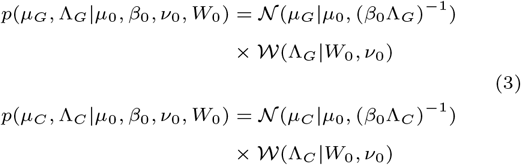

where 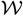 is the Wishart Distribution with *ν*_0_ as the degrees of freedom and *W*_0_ as the scale matrix.

We also set a zero mean normal distribution as *β_G_* and *β_C_* ‘s priors and a gamma distribution as the problem dependent *α_G_* and *α_C_* ‘s hyperpriors adapted from Macau [18]:

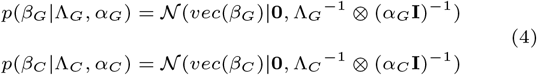

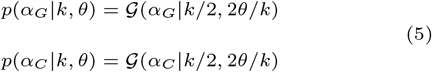

where *vec*(*β_X_*) is the vectorization of *β_X_*, ⊗ represents the Kronecker product and *α_X_* is the precision (*X* ∈ {*G, C*}). *k/*2 and 2*θ/k* are shape and scale, respectively. *k* and *θ* are hyperparameters which are set to 1.

### Gibbs sampler to impute dropout events

We use Markov Chain Monte Carlo (MCMC) algorithm to train Bfimpute, which is a sampling based approach to tackle the Bayesian inference problem. Bfimpute constructs a Markov Chain from a random initial value and after running the chain for 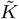 steps, it will eventually converge to its stationary distribution. Bfimpute then uses the average of 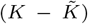 stationary stages to approximate the real distribution of *E* and gain the estimated values 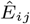 for dropouts:

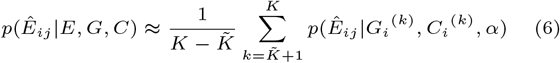

More specifically, Bfimpute chooses Gibbs sampler to achieve Bayesian matrix factorization. In every cycle, we sample the conditional distribution from the posterior distribution in Bayes’ theorem. Since the probabilistic models of genes and cells are symmetric, the conditional distributions over genes and the conditional distribution over cells have the same form. In particular, based on (1) and (2), the conditional probability for *G_i_* is:

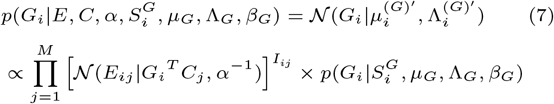

where

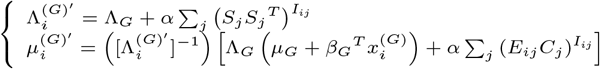

According to (2) and (3), we can derive the conditional probability for *μ_G_* and Λ_*G*_:

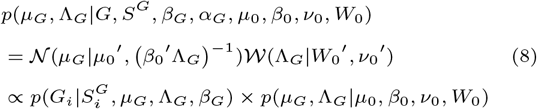

where

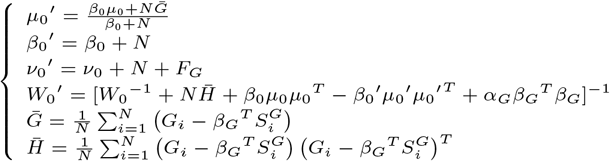

Considering (4) and (5), we get the conditional probability for *α_G_*:

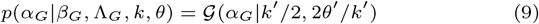

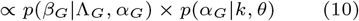

where

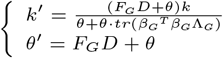

From (2) and (4), we are able to know the conditional probability for *β_G_*:

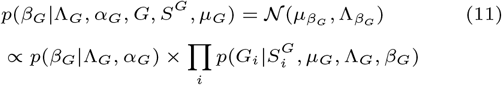

Because the size of the precision matrix 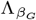 is too large to compute, we consider to do this part in an alternative way which is firstly proposed by Macau [18] by calculating:

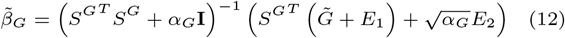

where 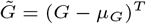, and each row of *E*_1_ ∈ ℝ^*N* ×*D*^ and 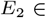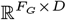 is sampled from 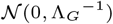.

**Algorithm 1.**
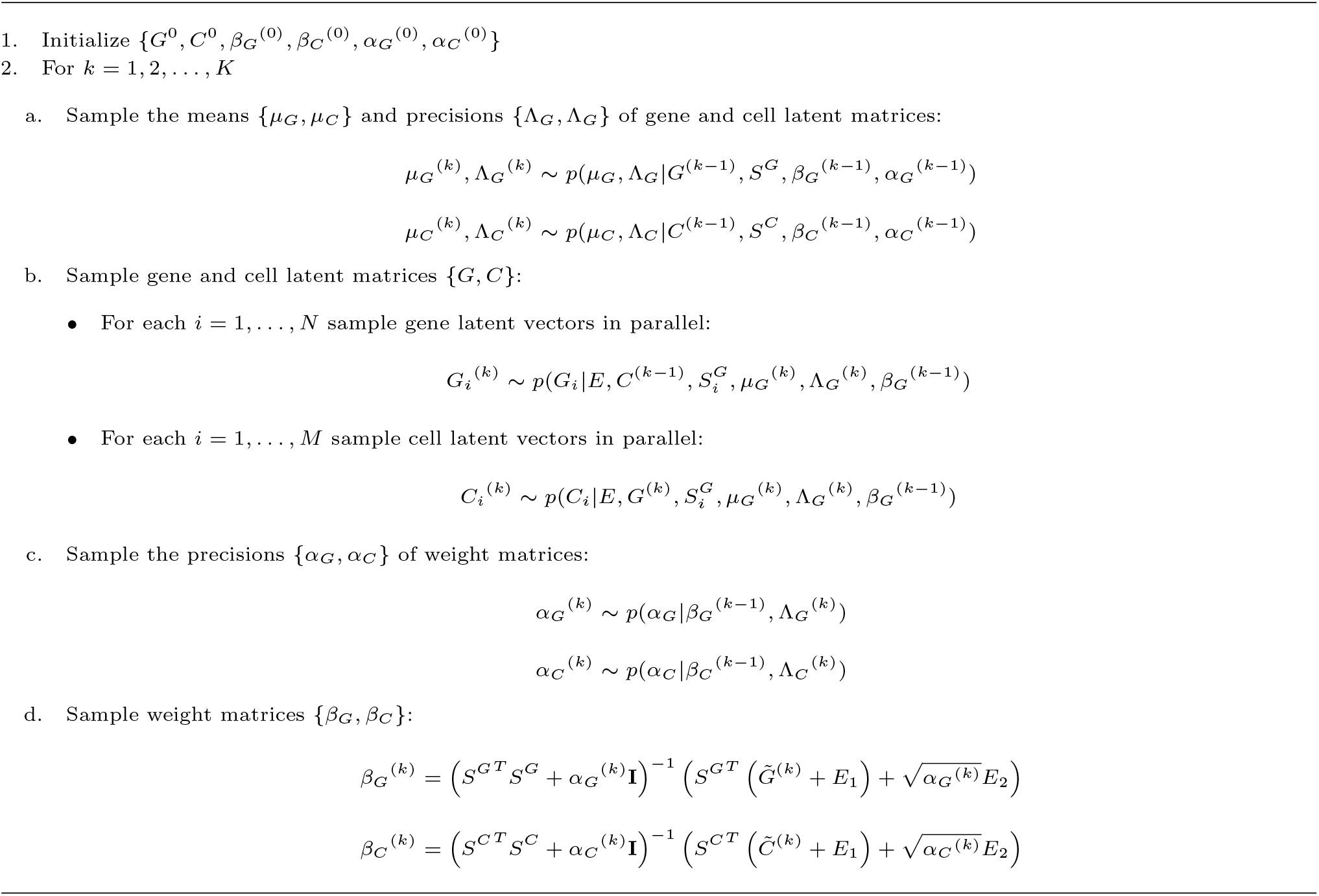
Gibbs sampling in Bfimpute

The Gibbs sampling steps of Bfimpute are shown in Algorithm 1:

### Generation of simulated data

We first simulated a single cell RNA-seq count matrix with 20000 genes and 500 cells evenly split into 5 groups using the scater(v1.16.2) [24] package and Splatter(v1.12.0) [25] package. The parameter which controls the probability that a gene will be selected as DE was set to 0.08 while the location and scale factor were set to 0.3 and 0.5, respectively. We used ‘experiment’ to add the global dropout for every cell. In order to show the universal applicability of Bfimpute, we further generated 6, 7, 8 groups of cells with 600, 700, 800 as total cell numbers and 10 runs for each data with different seeds using the same parameters mentioned above.

### Quality Control for real datasets

We did quality control (QC) (https://github.com/gongx030/ scDatasets) for all real datasets to ensure fairness for all methods before imputation except for PBMCs dataset (see details in Github). As the PBMCs dataset is based on 10X Genomics platform with an extremely high dropout rate, the QC step for PBMCs datasets could remove and lose nearly 80% genes.

### Evaluation metrics of clustering results

We used four evaluation methods: adjusted Rand index [26], Jaccard index [27], normalized mutual information (nmi) [28], and purity score, to analyse the agreement between true cluster labels and the spectral clustering [22] results on the first two Principle Components (PCs) of imputed matrix. Most of these four measurements vary from 0 to 1, with 1 indicating perfect match between them, except the adjusted Rand index which could yield negative values when agreement is less than expected by chance. The adjusted Rand index is an adjusted version of Rand’s statistic [29] which is the probability that a randomly selected pair is classified in agreement. The Jaccard index is similar to Rand Index, but disregards the pairs of elements that are in different clusters for both clusterings [30]. The normalized mutual information combines multiple clusterings into a single one without accessing the original features or algorithms that determine these clusterings. The purity score shows the rate of the total number of cells that are classified correctly.

## Results

We demonstrated the performance of Bfimpute in gene expression recovering, data visualization, cell subpopulation clustering, pseudotime and DE analysis on five publicly available scRNA-seq datasets (Supplementary Table 1), and we compared Bfimpute with six state-of-the-art imputation methods: scImpute, SAVER, VIPER, DrImpute, MAGIC, and SCRABBLE in the following sections.

### Bfimpute improves both visualization and cell type identification

PCA and t-distributed stochastic neighbor embedding (t-SNE) [31, 24] are two popular dimensionality reduction techniques often used to visualize high-dimensional scRNA-seq datasets. Since the dropout values were unknown in real datasets, we first tested accuracy of all different imputation methods using a simulated dataset where the ground truth was known. We applied the Splatter method to generate simulated datasets, which simulated many features observed in the scRNA-seq data, including zero-inflation, gene-wise dispersion, and differing sequencing depths between cells. To test the strength and robustness of different imputation methods, we simulated a wide range of datasets to include 5, 6, 7 and 8 different cell types (Methods section). Bfimpute achieved the most compact and well separated clusters on the simulation, followed by scImpute and DrImpute (Figure 2). For all different cell types simulations, we also evaluated the clustering performances by the evaluation metrics, where Bfimpute achieved the best scores for adjusted Rand index, Jaccard index, normalized mutual information and purity score compared to the raw data and other five imputation methods (Methods section).

**Fig. 2.**
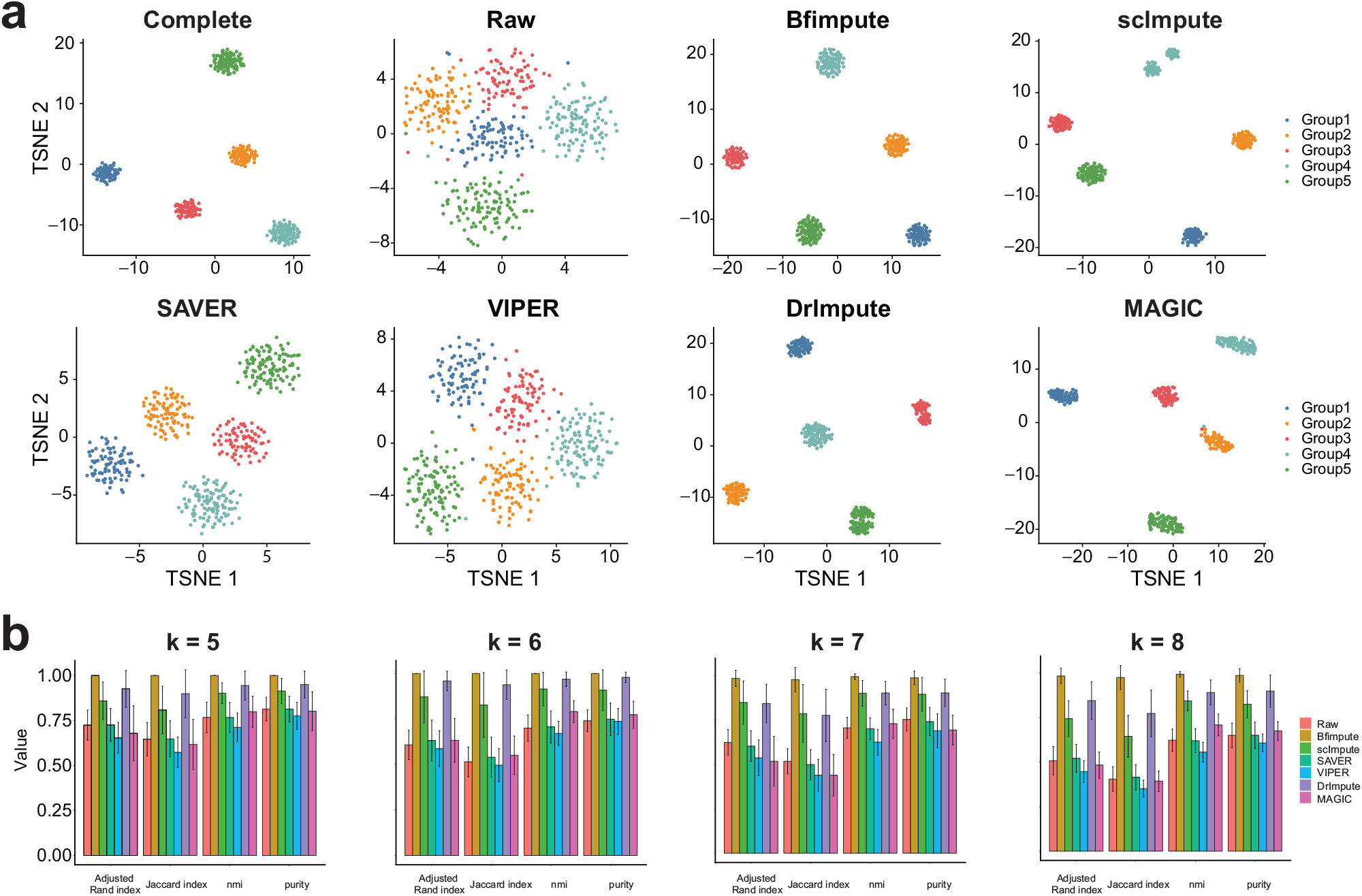
Bfimpute recovers dropout values and improves cell type identification in the simulated data. a. The scatter plots show the first two dimensions of the t-SNE results calculated from the complete data, the raw data, and the imputed data by Bfimpute, scImpute, SAVER, VIPER, DrImpute, and MAGIC. b. k represents the number of cell clusters in simuated data. The adjusted Rand index, Jaccard index, nmi, and purity scores of clustering results are based on the raw and imputed data.

We further used two real datasets for this analysis and the first two principal components (PCs) from PCA were plotted to compare every dataset across seven different conditions: raw dataset, and six imputed ones through the Bfimpute, scImpute, SAVER, VIPER, DrImpute, and MAGIC methods. We first applied all imputation methods to a real scRNA-seq dataset from a human embryonic stem (ES) cell differentiation study [2] to demonstrate the capacity of Bfimpute for improving the performance of data visualization. The dataset contains 1018 single cells from seven cell groups: Neuronal progenitor cells (NPCs), definitive endoderm cell (DEC), endothelial cells (ECs) and trophoblast-like cells (TBs) are progenitors differentiated from H1 human ES cells. H9 human ES cells and human foreskin fibroblasts (HFFs) were used as controls cells. The raw dataset (i.e. without imputation) clearly identified the cluster of HFF cells, however five other cell types were clustered very closely. After imputation by Bfimpute, the homogeneous subpopulations of H1 and H9 human ES cells were observed to substantially overlap and well separated from the rest of the progenitors. The DECs, ECs, HFFs, NPCs and TBs were also compactly clustered and well separated on the PCA plot (Figure 3a). Compared with the raw dataset, SAVER, VIPER and DrImpute had no significant improvement for cell groups identification. scImpute was the second best and generated similar compact cell groups as Bfimpute. We then compared clustering results of the spectral clustering algorithms [22] on the first two PCs to demonstrate the capability of Bfimpute to improve clustering accuracy in cell type identifications. For the true labels, we had seven cell types for this dataset, and we evaluated the clustering results by four different metrics: adjusted Rand index, Jaccard index, normalized mutual information (nmi), and purity (Methods section). All four metrics suggested Bfimpute achieved the best clustering accuracy compared with raw and other five imputation methods (Figure 3b). We also showed the comparison of visualization performance through t-SNE. t-SNE on the raw dataset can better identify the seven cell types comparing to PCA. Bfimpute, DrImpute and SAVER can further separate different cell groups and improve the visualization, however the other four imputation methods demonstrated worse t-SNE results than raw data (Supplementary Figure 1).

**Fig. 3.**
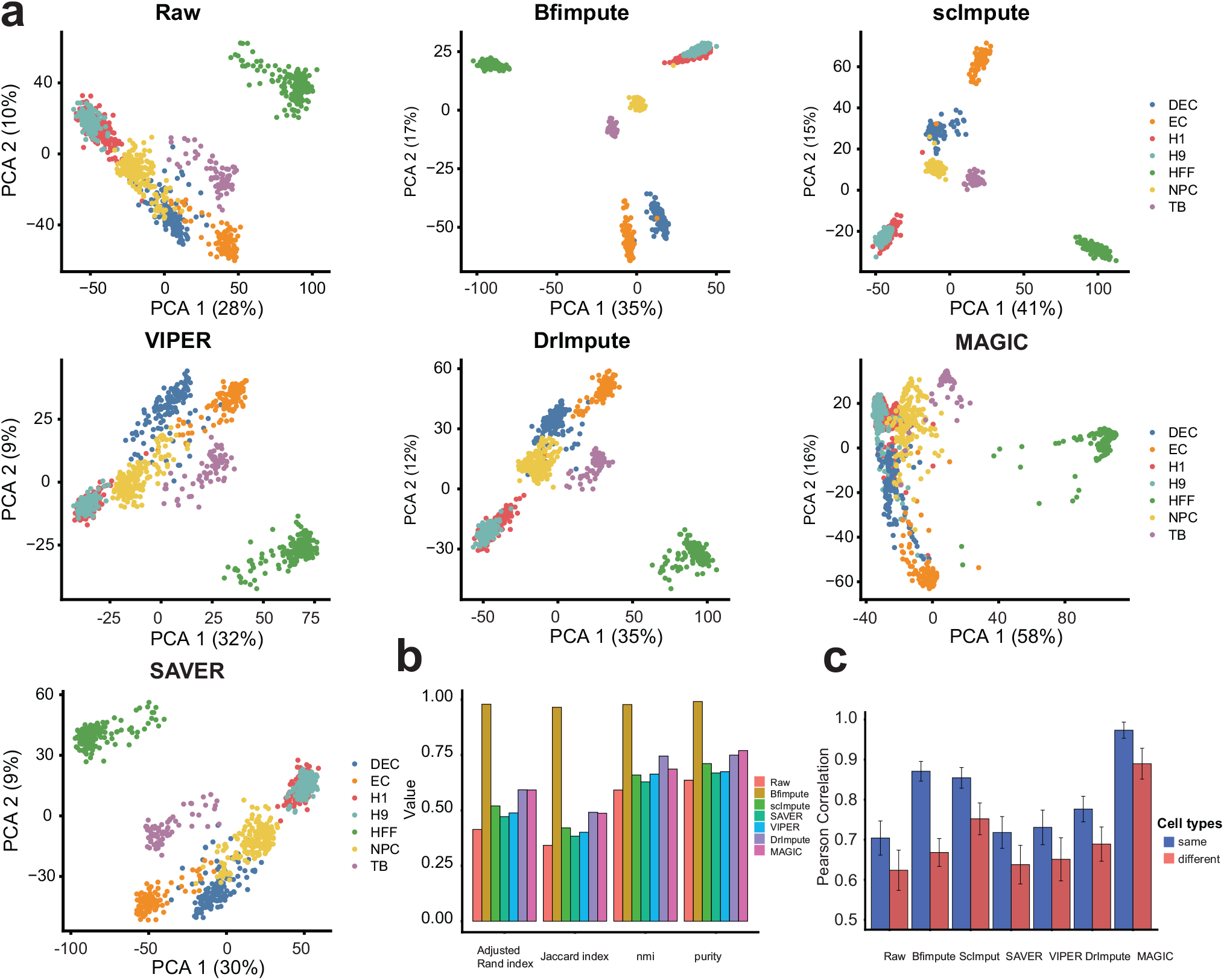
Bfimpute improves PCA visualization and cell type identification. a. The first two PCs calculated from the raw data, and the imputed data by Bfimpute, scImpute, VIPER, DrImpute, MAGIC, and SAVER. b. The adjusted Rand index, Jaccard index, nmi, and purity scores of clustering results based on the raw and imputed data. c. Average Pearson correlations between any two cells from same type and different type.

To illustrate the recovering of dropouts in individual cells by imputation, we calculated the Pearson correlation from *log*_10_-transformed read counts between every pair of cells in the same type and from different cell types. This result indicated imputation did recover the zero counts in every cell and the Pearson correlation increased from 0.70 to 0.87 for Bfimpute, 0.85 for scImpute, 0.72 for SAVER, 0.73 for VIPER, 0.78 for DrImpute, and 0.97 for MAGIC (Figure 3c, blue bars). One scatter plot of correlations between two randomly selected stem cells of the same cell type was demonstrated in Supplementary Figure 2. As we expected, imputation methods usually increased the Pearson correlation between any two cells in the same cell type. Imputation should not increase the correlation between cells in different cell types by disregarding the biological variation between them. Among all imputation methods, MAGIC achieved the highest correlation in the same cell type, but the correlation between different cell types was also the highest (Figure 3c, red bars). Bfimpute demonstrated the best balance, by maximizing the difference between correlation for the same over different cell types.

We further investigated Bfimpute’s performance of visualization and cell type identification on another zebrafish [3] scRNA-seq dataset. This dataset contains 246 single cells from six cell groups, and Hematopoietic stem and progenitor cells (HSPCs) and HSPCs/thrombocytes among them come from one defined cell type with expected heterogeneity. After the QC step, the zebrafish dataset was still sparse with zeros composing over 87.5% of the total counts. The comparison of visualization performance via PCA on the raw and six imputed datasets is shown in Supplementary Figure 3. The raw dataset only roughly identified the cluster for neutrophil cells, whereas cells from other cell types were mixed and spread wildly. After imputation by Bfimpute, four distinct immune cell subpopulations can be identified for neutrophils, T, Natural Killer (NK) and B cells, where the cluster members were much more compact compared to those of the raw dataset. Neutrophils, T, NK and B cells were distantly positioned on the PCA plot. HSPCs and HSPCs/thrombocytes were from one defined cell type with expected heterogeneity, so after Bfimpute’s imputation, they were still spatially closer than other cells (Supplementary Figure 3a). The raw data and the imputed data by other five imputation methods did not correctly identify the four immune cell subpopulations. Clustering accuracy results from the four metrics for Bfimpute were better than the other five imputation methods, and Bfimpute achieved a better correlation for the same cell type without loosing variation between different cells types (Supplementary Figure 3b,c).

### Bfimpute improves DE and pseudotime analysis

DE analysis is widely used in bulk RNA-seq data. Performing DE analysis for scRNA-seq data to reveal the stochastic nature of gene expression in single cells is challenging since scRNA-seq data suffers from high dropout events. However, it has been proven that good imputation methods could lead to a better agreement between scRNA-seq and bulk RNA-seq data of the same biological condition on genes known to have little cell-to-cell heterogeneity. We utilized a real dataset by Chu et al [2] with both bulk and scRNA-seq data available on human embryonic stem cells and definitive endoderm cells (DEC) [32, 33], to compare Bfimpute with the raw dataset and other five imputation methods for DE analysis. This dataset contained six samples of bulk RNA-seq (four in H1 ES cells and two in DEC) and 350 samples of scRNA-seq (212 in H1 ES cells and 138 in DEC). The percentages of zero entries were 8.8% in bulk data and 44.9% in scRNA-seq data, respectively. We first performed DE analysis in the bulk data and identified the top 200 DE genes by DESeq2 [10]. We then plotted these 200 genes’ expression profiles in scRNA-seq data for seven conditions: raw dataset, Bfimpute, scImpute, SAVER, VIPER, DrImpute, and MAGIC. We found these top 200 genes’ expression profiles after Bfimpute’s imputation demonstrated better concordance with those in bulk data (Figure 4a). To further evaluate whether imputation improves DE analysis in scRNA-seq data, we first used DESeq2 to identify DE genes for raw scRNA-seq dataset and scRNA-seq datasets after six different imputations. We then generated different lists of DE genes for the bulk data by applying different thresholds for false discovery rates of genes. Finally for every threshold, we compared the DE genes for the bulk data and scRNA-seq data of those seven different conditions and calculated the AUC values for each condition. The AUC values suggested all imputation methods improved DE analysis. Bfimpute generated DE genes most consistent with the bulk data (AUC values raw: 0.568, Bfimpute: 0.670, scImpute: 0.665, SAVER: 0.624, VIPER: 0.639, DrImpute: 0.657 and MAGIC: 0.668).

**Fig. 4.**
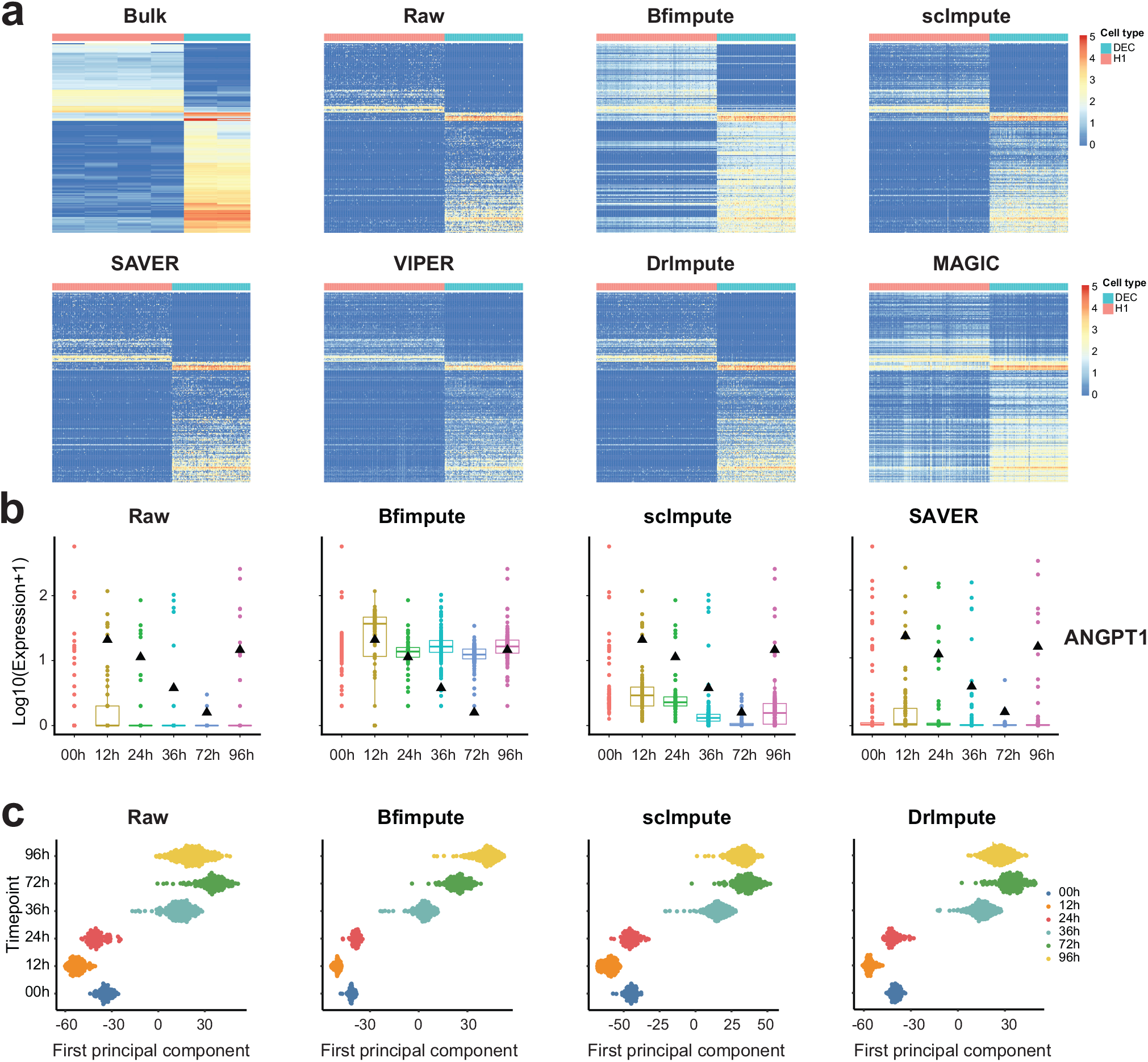
Bfimpute improves DE and pseudotime analysis. a. The expression profiles of the top 200 DE genes detected in the bulk data by DESeq2 for seven conditions: raw dataset, Bfimpute, scImpute, SAVER, VIPER, DrImpute, and MAGIC. b. Time-course expression patterns of the example gene ANGPT1 that is annotated with GO term “endoderm development”. The small black triangles marks the average bulk data for each time point. c. The first principal component is plotted to show cells of different time points along the differentiation.

Bulk data for the same biological condition was provided and could be used as a gold standard to compare the average gene expression level with the scRNA-seq data, even though the scRNA-seq data presented more cell-to-cell variation. We expected that average gene expression level in the scRNA-seq data was highly correlated with bulk RNA-seq data. To investigate this, we plotted correlations between gene expression in single-cell and bulk data and found that all imputation methods did improve the correlation between bulk and scRNA-seq data, and Bfimpute, MAGIC and scImpute had the best improvement (Supplementary Figure 4). We further selected several genes (e.g., ANGPT1,GDF3, BMP4, EPB41L5) of DECs from different time points to plot their average gene expression levels in both bulk and scRNA-seq data. These genes were annotated with the GO term “endoderm development”, and they were likely to be affected by dropout events [13, 34]. Imputed read counts for these genes by Bfimpute showed higher gene expression correlation and better consistency with the bulk data (Figure 4b and Supplementary Figure 5, 6).

In addition to the DE analysis, we also used the time course scRNA-seq data [2] from the same Chu et al study to show Bfimpute improved gene expressions temporal dynamics through pseudotime analysis. In this dataset, a total of 758 single cells were captured and profiled by scRNA-seq at 0, 12, 24, 36, 72, and 96 h of differentiation. We first applied Bfimpute, scImpute and Drimpute to the raw scRNA-seq data with true cell type labels, and then study how the time-course expression patterns change in the imputed data. The PCA results showed that imputed read counts by Bfimpute better distinguished cells of different time points and the six time points cell groups were compact (Supplementary Figure 7a), and the first principle component from PCA indicated that imputed read counts from Bfimpute reflected more accurate transcriptome dynamics along the different time course (Figure 4d). Bfimpute could better differentiate the last two time points (72h and 96h). In the next section, we will discuss impuation with the aid of cell type labels more in details.

### Bfimpute improves performance with the aid of additional experimental information

Imputation methods including Bfimpute, scImpute and DrImpute all first identified similar cells based on clustering, and imputation was then performed by leveraging the expression values from similar cells. Being able to first identify the appropriate cell groups enhanced the ability of imputing the dropout events. A substantial number of scRNA-seq studies have identified cell types from experimental design or marker genes. We applied Bfimpute, scImpute and DrImpute to the raw scRNA-seq data with true cell type labels in three real datasets we have used before, and two more new real datasets. In this study, SAVER, VIPER, DrImpute, and MAGIC were excluded since they were not applicable to use cell labels. We then investigated again the PCA and t-SNE visualizations for cell subpopulations identification. Our results showed Bfimpute outperformed the other two methods and clearly differentiated almost every cell group in different datasets (Figure 5 and Supplementary Figure 7, 8). For the human embryonic stem cell dataset, Bfimpute further correctly identified three outlier cells into correct groups compared to the previous imputation without cell labels (see Figure 5a versus Figure 3a: one EC (orange point), one DEC (blue point), and one NPC (yellow point) cell were brought back to the corresponding EC, DEC and NPC cell groups, respectively). H9 cells were also further apart from H1 cells in the vertical dimension. For the zebrafish dataset, even the most mixed B, NK, T cells (blue, green, and yellow colors) from the raw dataset were separated from each other after Bfimpute’s imputation, and HSPCs and HSPCs/thrombocytes cells were spatially close, but split into two cell groups (Figure 5b and supplementary Figure 7b).

**Fig. 5.**
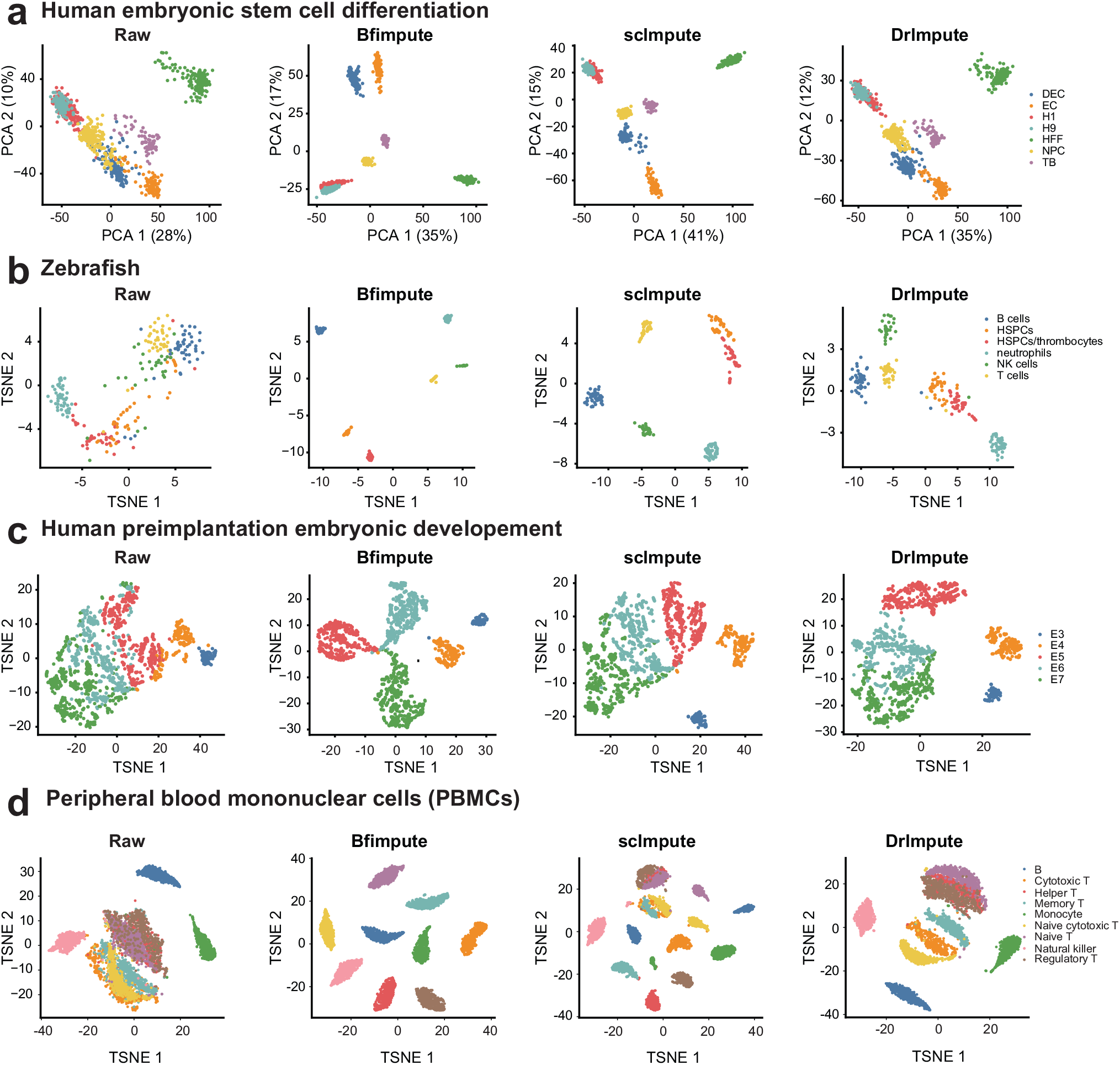
Bfimpute with labels improves PCA and t-SNE visualizations and cell type identification. a. The first two PCs calculated from the raw data, and the imputed data by Bfimpute, scImpute, and DrImpute for the human embryonic stem cell differentiation study. b. The first two dimensions from the raw data, and the imputed data by Bfimpute, scImpute, and DrImpute for the zebrafish data. c. The first two dimensions from the raw data, and the imputed data by Bfimpute, scImpute, and DrImpute for the human preimplantation embryonic development. d. The first principal component is plotted to show cells of different time points along the embryonic development.

To test Bfimpute with another kind of cell-label information, we used a human preimplantation embryonic development dataset (t-SNE and pseudotime analyses are shown in Figure 5c). The Petropoulos dataset [4] included single cells from five stages of human preimplantation embryonic development, ranging from developmental day (E) 3 to 7. The five different stages were clearly distinguished from each other after Bfimpute’s imputation.

We also applied three imputation methods to a large 10X dataset generated by the high-throughput droplet-based system. To generate this dataset, we randomly selected 500 cells from nine immune cell types, so it contained a total of 4500 peripheral blood mononuclear cells (PBMCs) [12, 5]. In the raw data, 98.3% read counts are exactly zeros. Our PCA and t-SNE results indicated that Bfimpute’s imputation identified nine immune cell types from raw data (5d). In summary, these results suggested that Bfimpute with the aid of labels always further improved visualization and identification of cell subpopulations, and the downstream analysis.

SCRABBLE is another recent approach integrating bulk data to impute dropout events in scRNA-seq data. Since Bfimpute can easily adopt bulk data as additional information into the gene latent matrix, we have also tested if bulk data can further improve performance. In the scRNA-seq dataset of human embryonic stem cells with bulk data, we did not observe significant differences between Bfimpute and Bfimpute with bulk data as additional information (Supplementary Figure 8 versus Figure 5a). The reason could be that similar gene level information has less effect than similar cell level information for the imputation of dropout events. We also found that in these scRNA-seq datasets, SCRABLE’s performance after integrating cell labels information with bulk data, was not better than Bfimpute (Supplementary Figure 8).

## Discussion and Conclusion

ScRNA-seq has become an indispensable tool in recent years, as it has made it possible to study genome-wide transcriptomes in single cell resolution. Due to sequencing technical issues, a large proportion of dropout events exist in scRNA-seq data, which limit its usefulness. Several approaches have been proposed to solve this problem, with modest results. In this study, we introduced Bfimpute to recover dropout events in scRNA-seq data. We have shown that Bfimpute can improve performance in recovering gene expression detected by bulk RNA-seq, as well as in downstream analyses, including identification of cell sub-populations, differential expressed genes and gene expressions temporal dynamics.

Bfimpute uses a fully Bayesian probabilistic matrix factorization by substituting hyperparameters with hyperpriors and performing Gibbs sampling for the approximate inference. The advantage of this Bayesian model is that it provides a predictive distribution instead of just a single number during recovering each dropout event, and the confidence in the prediction can be quantified and considered into the model. The use of a full Bayesian model proved to be a considerable advantage for Bfimpute to outperform other imputation methods.

Bfimpute imputes two latent cell and gene matrices for each cell group through a Gibbs sampling process, and reaches a stationary state to generate the final cell-gene expression matrix, in which the dropout events will be recovered. Another advantage of Bfimpute is being able to integrate any gene or cell related information of scRNA-seq data into these two latent gene and cell matrices to impute missing values. Information from both similar cells or/and bulk data can be easily integrated into our model. Even though scImpute and DrImpute have a similar functionality in this respect, that allows them to impute dropout events with the aid of number of cell types or cell labels, they fail to achieve as good performance as Bfimpute for most of scRNA-seq data that we tested. Any resource provided by the users from the cell level and gene level could be used as additional information to improve dropout events imputation in scRNA-seq data in the future.

## Key Points

- Imputation to recover dropout events for scRNA-seq data is important for determining genome-wide transcriptomes in single cell resolution.
- Bfimpute uses a fully Bayesian probabilistic matrix factorization by substituting hyperparameters with hyperpriors and performing Gibbs sampling for approximate inference.
- The advantage of this Bayesian model is that it provides a predictive distribution instead of just a single number during recovering each dropout event, and the confidence in the prediction can be quantified and considered into the model.
- Bfimpute is able to integrate any gene or cell related information of scRNA-seq data into these two latent gene and cell matrices to impute missing values.
- Bfimpute achieves better accuracy than other six widely used scRNA-seq imputation methods on simulated and real scRNA-seq data, as measured by several different evaluation metrics.

## Supporting information

Bfimpute_SupplementaryInformation

## Author contributions

X.Z. conceived and led this work. Z.H.W. and X.Z. designed the model and implemented the Bfimpute software. Z.H.W., J.L.L, W.S and X.Z led the data analysis. Z.H.W, W.S and X.Z wrote the paper with feedback from J.L.L and L.Z.

