## Supplementary material for "Bfimpute: A Bayesian factorization method to recover single-cell RNA sequencing data": Bfimpute_SupplementaryInformation

**Wen et al.**

**Supplementary Information**

### Supplementary Tables and Figures

| Datasets | #Clusters | Cells | Data missing rate | Cell label definition | Ref. |
|---|---|---|---|---|---|
| Chu et al. | 7 | 1018 | 45.46281% | Human embryonic stem cell differentiation | 2 |
| Chu et al. | 6 | 758 | 48.39046% | Human embryonic stem cell differentiation (time course data) | 2 |
| Tang et al. | 6 | 246 | 87.5279% | Zebrafish | 3 |
| Petropoulos et al. | 5 | 1529 | 55.61331% | Stages of human preimplantation development | 4 |
| Zheng et al. | 9 | 4500 | 98.2839% (without QC) | Peripheral blood mononuclear cells | 5 |

Supplementary Table 1. Clusters k represents the number of cell clusters reported in the original study.

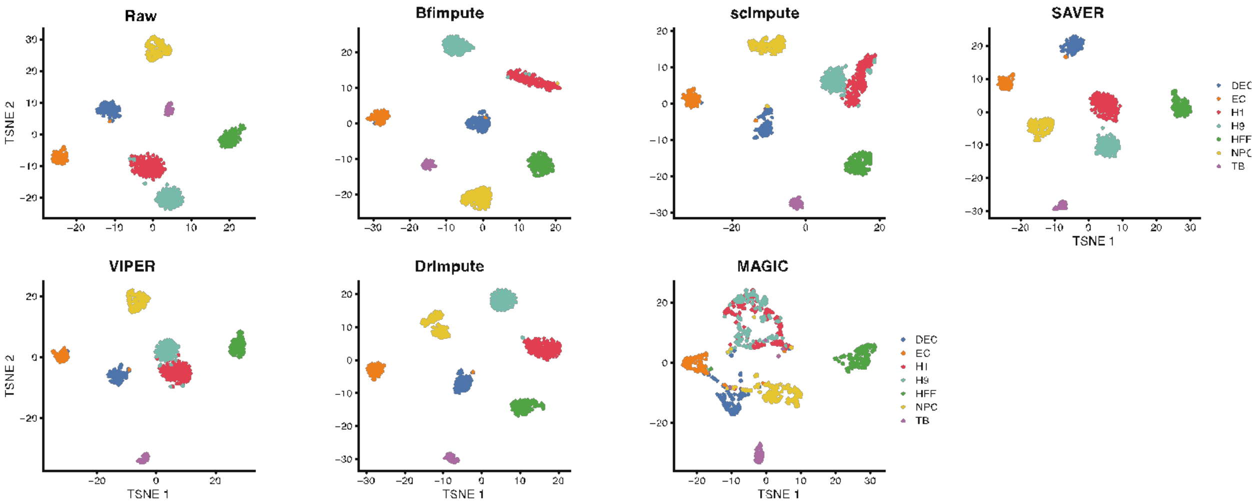


Supplementary Figure 1. The scatter plots show the first two dimensions of the t-SNE results calculated from the complete data, the raw data, and the imputed data by Bfimpute, scImpute, SAVER, VIPER, DrImpute, and MAGIC for Human embryonic stem cell differentiation study.

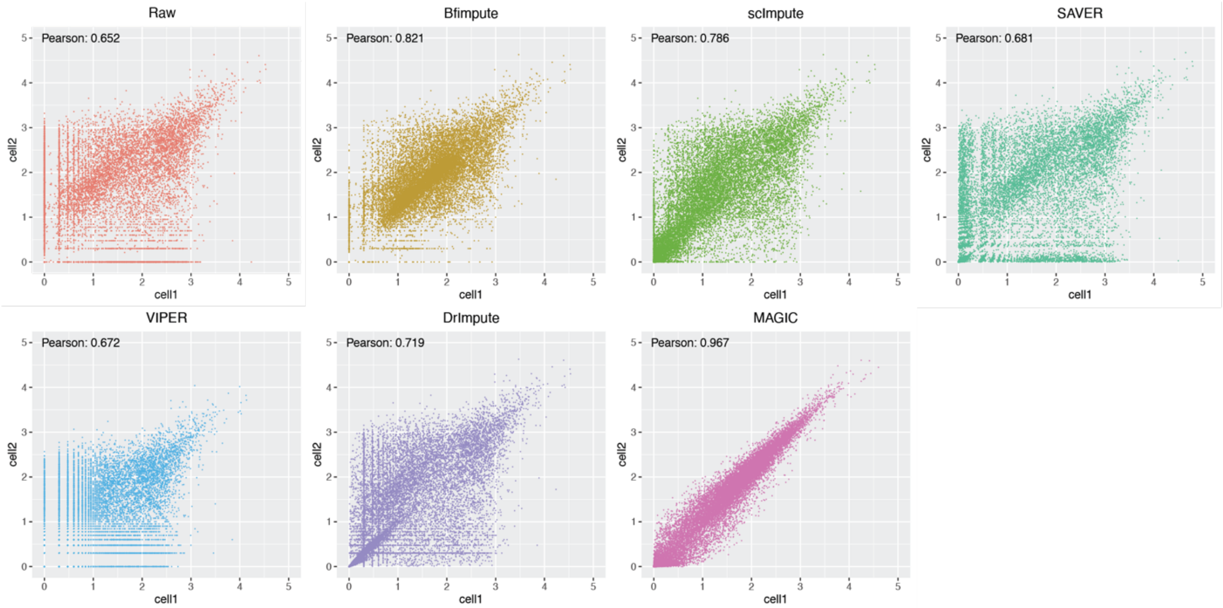


Supplementary Figure 2. The scatter plots the raw and imputed gene expression levels of two randomly selected stem cells of the same cell type. Pearson correlation coefficients between the two cells are shown on the top-left of each scatter plot.

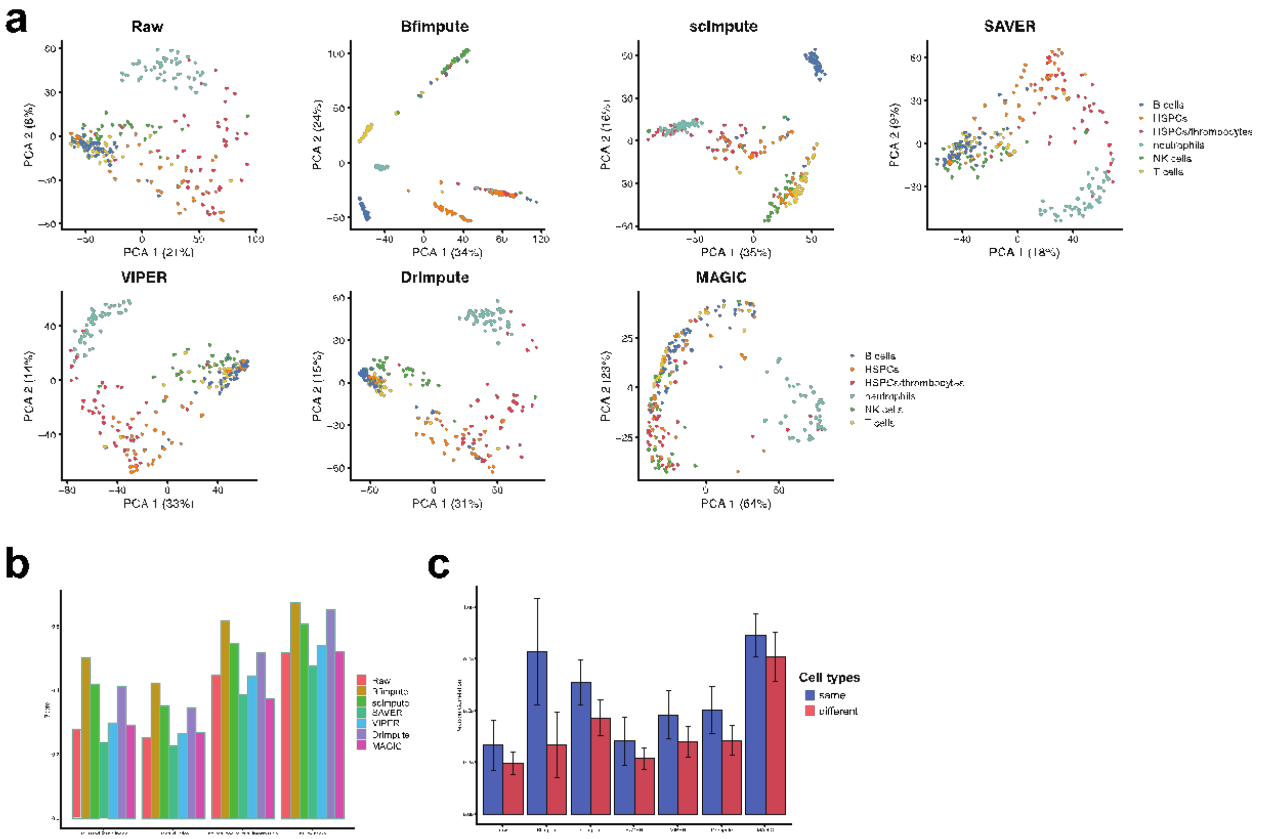


Supplementary Figure 3. Bfimpute improves PCA visualization and cell type identification for Zebrafish data. a. The first two PCs calculated from the raw data, and the imputed data by Bfimpute, scImpute, VIPER, DrImpute, MAGIC, and SAVER. b. The adjusted Rand index, Jaccard index, nmi, and purity scores of clustering results based on the raw and imputed data. c. Average Pearson correlations between any two cells from same type and different type.

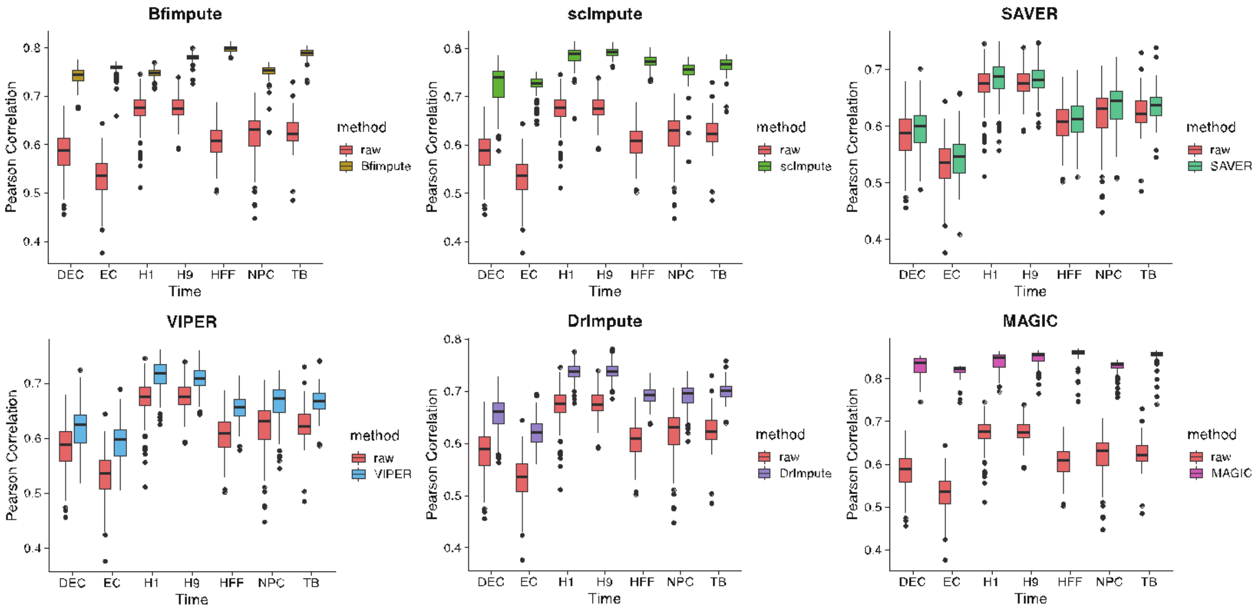


Supplementary Figure 4. Pearson correlations between gene expression levels in single-cell and bulk data. The Pearson correlation coefficients are calculated between each cell and averaged bulk data of each cell type for both raw and imputed data. The Pearson correlations for imputed data are significantly higher than those for raw data.

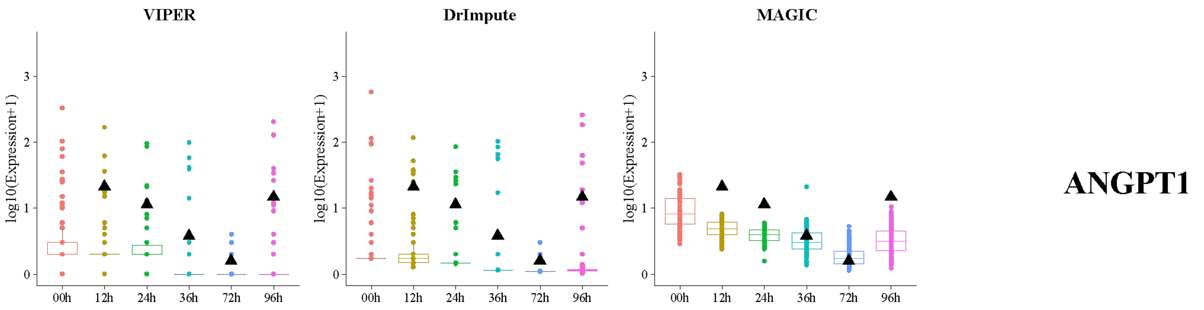


Supplementary Figure 5. Time-course expression patterns of the example gene ANGPT1 that is annotated with GO term "endoderm development". The small black triangles mark the average bulk data for each time point.

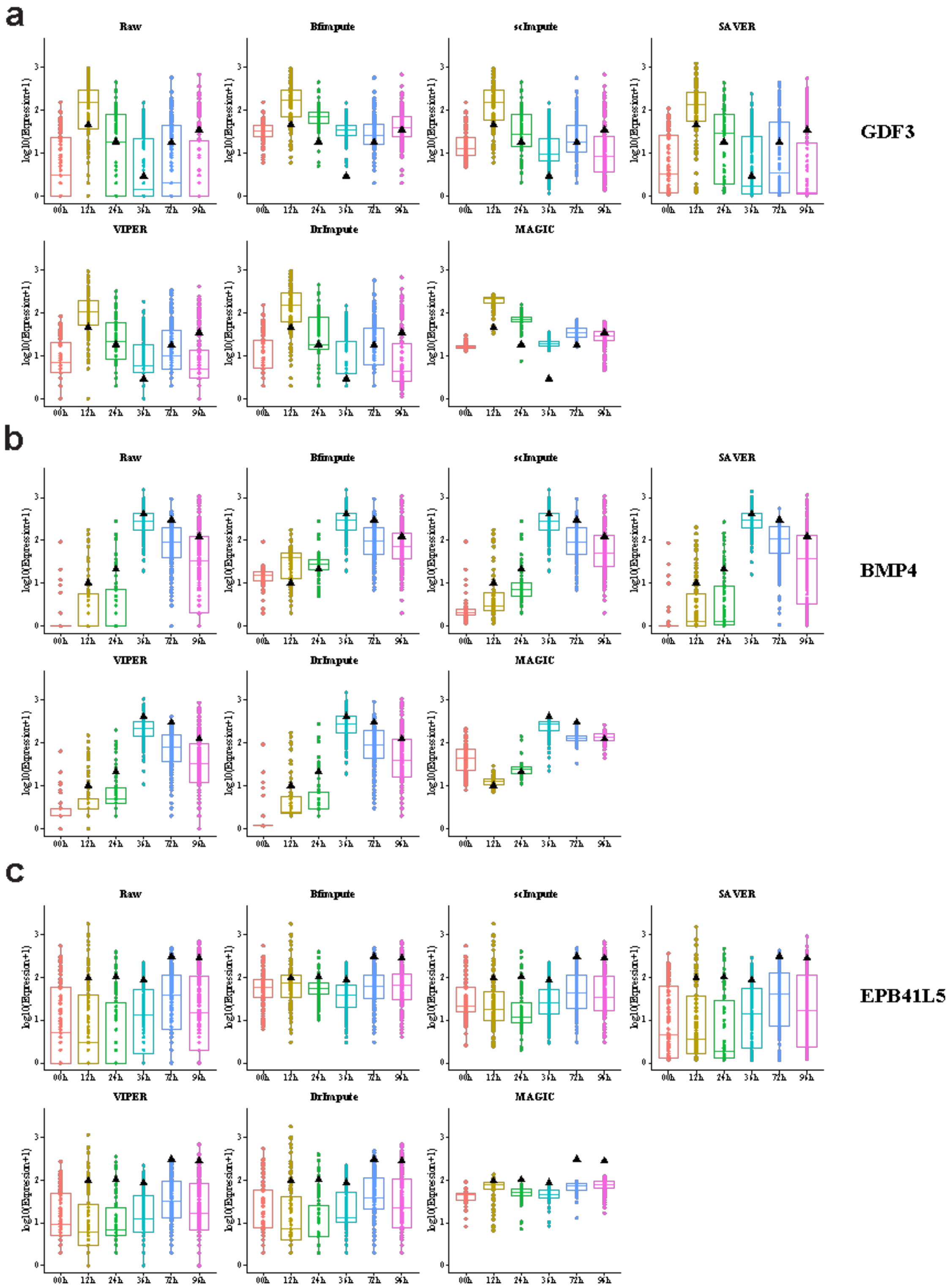


Supplementary Figure 6. Time-course expression patterns of the three more example genes that are annotated with GO term "endoderm development". The small black triangles mark the average bulk data for each time point. Imputed data by Bfimpute is highly correlated with the bulk dat

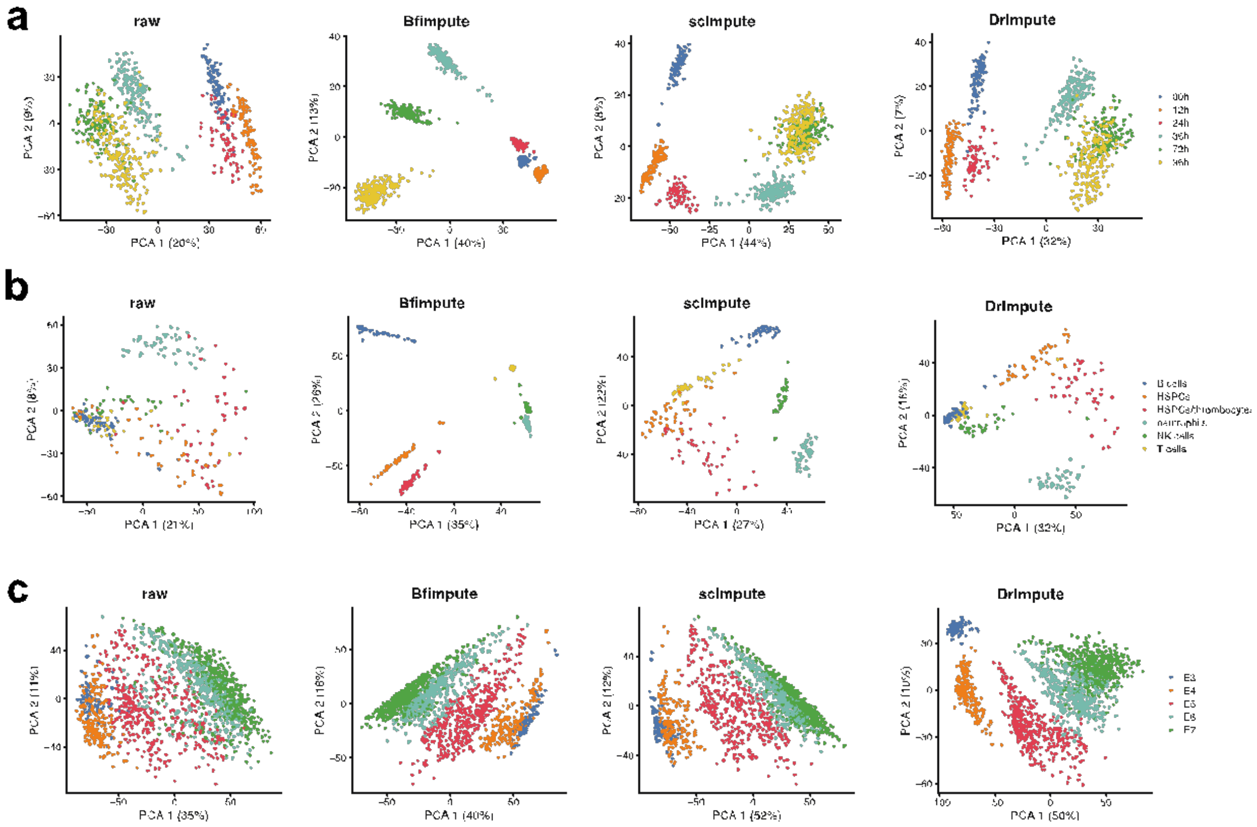


Supplementary Figure 7. Bfimpute with labels further improves PCA visualizations. a. The first two PCs calculated from the raw data, and the imputed data by Bfimpute, scImpute, and DrImpute for the human embryonic stem cell differentiation time course study. b. The first two PCs calculated from the raw data, and the imputed data by Bfimpute, scImpute, and DrImpute for the Zebrafish data. c. The first two PCs calculated from the raw data, and the imputed data by Bfimpute, scImpute, and DrImpute for the human preimplantation embryonic development study.

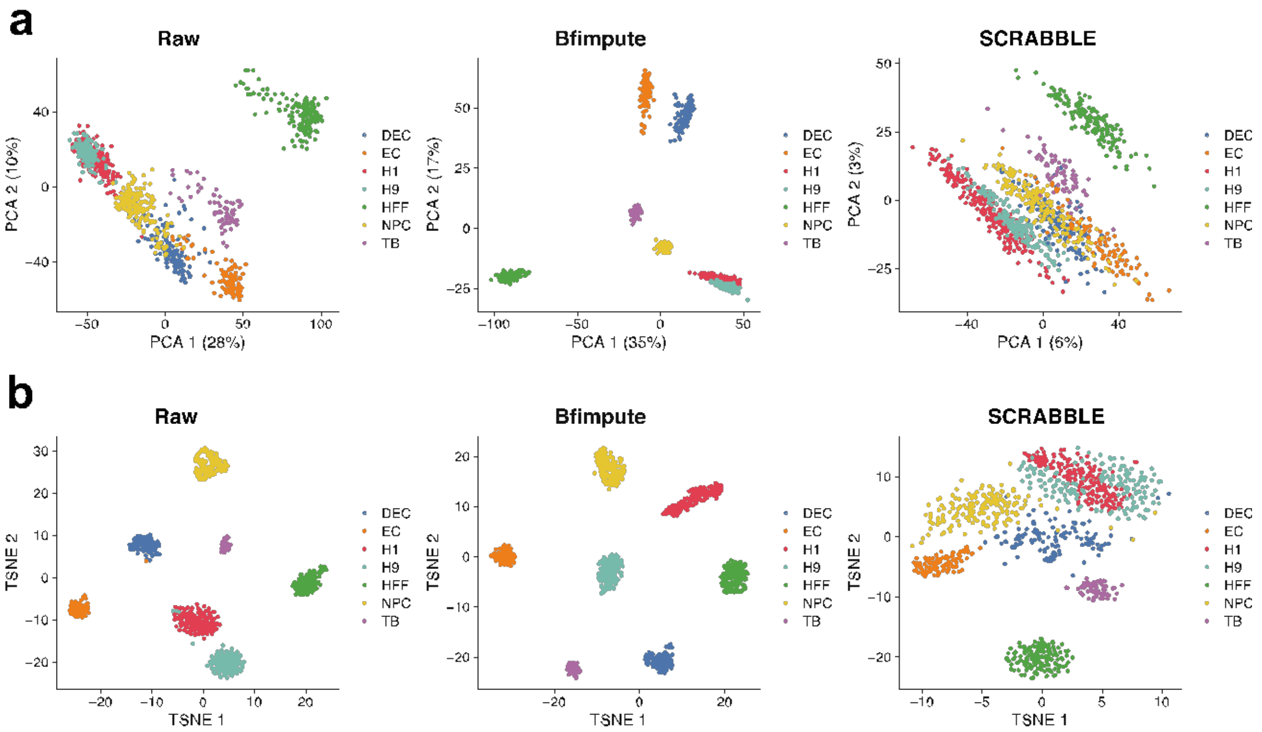


Supplementary Figure 8. Imputation with Bulk data. a. The first two PCs calculated from the raw data, and the imputed data by Bfimpute and SCRABBLE for Human embryonic stem cell differentiation study. b. The scatter plots show the first two dimensions of the t-SNE results calculated from the raw data, and the imputed data by Bfimpute and SCRRABLE for Human embryonic stem cell differentiation study.
